# Polygenic sex determination produces modular sex polymorphism in an African cichlid fish

**DOI:** 10.1101/2021.10.09.463756

**Authors:** Emily C. Moore, Patrick J. Ciccotto, Erin N. Peterson, Melissa S. Lamm, R. Craig Albertson, Reade B. Roberts

**Author notes:** ECM, RBR, and RCA conceived of the experiments, ECM took all photographs, performed all behavior experiments, and hormone assays, ECM and ENP performed gonad histology and sex genotyping, MSL performed gut length experiment, PJC and RCA analyzed head and body morphometrics, ECM performed all other statistical analyses, ECM and RBR wrote the manuscript with feedback from all authors. The authors declare no competing interests. To whom correspondence should be addressed. ECM,; RBR.

## Abstract

For many vertebrates, a single genetic locus initiates a cascade of developmental sex differences in the gonad and throughout the organism, resulting in adults with two, phenotypically distinct sexes. Species with polygenic sex determination (PSD) have multiple interacting sex determination alleles segregating within a single species, allowing for more than two genotypic sexes, and scenarios where sex genotype at a given locus can be decoupled from gonadal sex. Here we investigate the effects of PSD on secondary sexual characteristics in the cichlid fish *Metriaclima mbenjii*, where one female (W) and one male (Y) sex determination allele interact to produce siblings with four possible sex classes: ZZXX females, ZWXX females, ZWXY females, and ZZXY males. We find that PSD in *M. mbenjii* produces an interplay of sex-linkage and sex-limitation resulting in modular variation in morphological and behavioral traits. Further, the evolution or introgression of a novel sex determiner creates additional axes of phenotypic variation for varied traits, including genital morphology, craniofacial morphology, gastrointestinal morphology, and home tank behaviors. In contrast to single-locus sex determination, which broadly results in sexual dimorphism, polygenic sex determination can induce higher-order sexual polymorphism. The modularity of secondary sexual characteristics produced by PSD provides novel context for understanding the evolutionary causes and consequences of maintenance, gain, or loss of sex determination alleles in populations.

**Significance Statement:** Sex differences in traits can occur when those traits are modified by genetic factors inherited on sex chromosomes. We investigated how sex differences emerge in a species with more than one set of sex chromosomes, measuring a variety of morphological, physiological, and behavioral traits. Rather than exhibiting sexual dimorphism associated with primary sex, the species has higher-order sexual polymorphism in secondary sexual characteristics, or more than two phenotypic sexes. Variation in secondary sexual characteristics is modular, involving the interplay of sex-linked and sex-limited traits. Our findings provide novel implications for how sex determination systems and whole-organism fitness traits co-evolve, including that significant creation or loss of variation in diverse traits can occur during transitions among sex chromosome systems.

---

In sexually reproducing organisms, divergent optima for sexes (1) and sex-specific reproductive signaling (2) result in the evolution of sex differences in phenotype. It can be difficult to disentangle whether traits that differ by sexes are influenced by the sex determination gene itself (pleiotropy), driven by sex-specific hormonal changes during development (sex-limited), or influenced by variation in genes linked to the sex determining gene (sex-linkage) (3). In some taxa, studies have found weaker-than-expected evidence for sex-linkage underlying sexually dimorphic traits (3, 4). In others, there is clear evidence of linkage between a sex determiner and a trait undergoing sexually-antagonistic selection (5). Whether sexual dimorphism is related to mate choice or differential life history strategies, the mechanisms underlying these differences can provide potent evolutionary constraint from sexual conflict (6) and additional opportunity once that sexual conflict is resolved (5).

Despite the importance of sex determination in the maintenance of distinct, reproductively compatible sexes, sex determination systems are variable, and there have been many evolutionary transitions between genetic sex determination systems in vertebrates (7–9). In most single-locus genetic sex determination systems, sex is determined through inheritance of a dominant allele from the parent of the same sex—in ZW systems, mothers pass on a W to daughters and a Z to sons; in XY systems, fathers pass on a Y to sons and X to daughters. Polygenic sex determination (PSD) systems are more complex, involving segregation of more than one genetic sex determiner in a single population, and are found in both animal and plant taxa (10–12). Where multiple sex determiners are present in a single individual, their epistatic interactions determine primary sex, often in a hierarchical fashion (10, 13). Importantly, the loci involved in PSD provide multiple sites in the genome for sex linkage, and interactions which have the potential to increase variation in sex-limited gene expression.

The cichlid fishes of East Africa are an excellent model to understand the evolution of sexual polymorphism as they have undergone a recent adaptive radiation characterized by diversification in numerous phenotypes that also display robust sex differences, including pigmentation, behavior, and morphology (14–16). Additional diversity is found in sex determination systems, with at least five different genetic sex determination loci described in the Lake Malawi cichlid flock, co-segregating in some species to produce PSD systems (17).

Quantitative genetic mapping of sex-influenced traits such as pigmentation (5, 13, 18) and craniofacial morphology (19) has supported an association with these traits and autosomal sex loci in cichlids, though it is difficult to resolve sex-linkage from sex-limitation from these studies. In the cichlid system, sex-associated variation has been demonstrated in trophic traits including gill rakers (20) and host-gut-microbiota interactions (21), and likely reflects adaptations and sexual conflicts related to mouthbrooding and unequal parental investment. Differences in behavioral strategies within and between sexes have been well documented in cichlids, particularly related to male-male behavioral competitions influencing mating success (22–24). However, links between behavior and mechanism made thus far involve socially mediated modulation of sex steroid hormones and little is known about the genetic basis of behavioral variation. One possible exception is the finding that sex-linked blotch color morph females in Lake Victoria are more aggressive than non-blotched females of the same species, suggesting behavioral variation is associated with which sex chromosomes individuals possess (25).

In this study, we measured morphological, physiological, and behavioral phenotypes in a Lake Malawi cichlid fish species with PSD, *Metriaclima mbenjii*. This species has a multilocus sex determination system, as previously described for its congener *M. pyrsonotos* (11). In these species, a putatively-ancestral XY sex determination system on chromosome 7 (chr. 7) interacts epistatically with a ZW system on chromosome 5 (chr. 5), with individuals inheriting both the Y and the W allele developing as female ((11); Fig.1A). The W sex determination allele is tightly linked to an allele of pax7a producing a distinct blotched color morph only in females carrying the W; the blotching putatively provides an alternative form of crypsis to plain brown female coloration (5, 26)

**Fig. 1.**
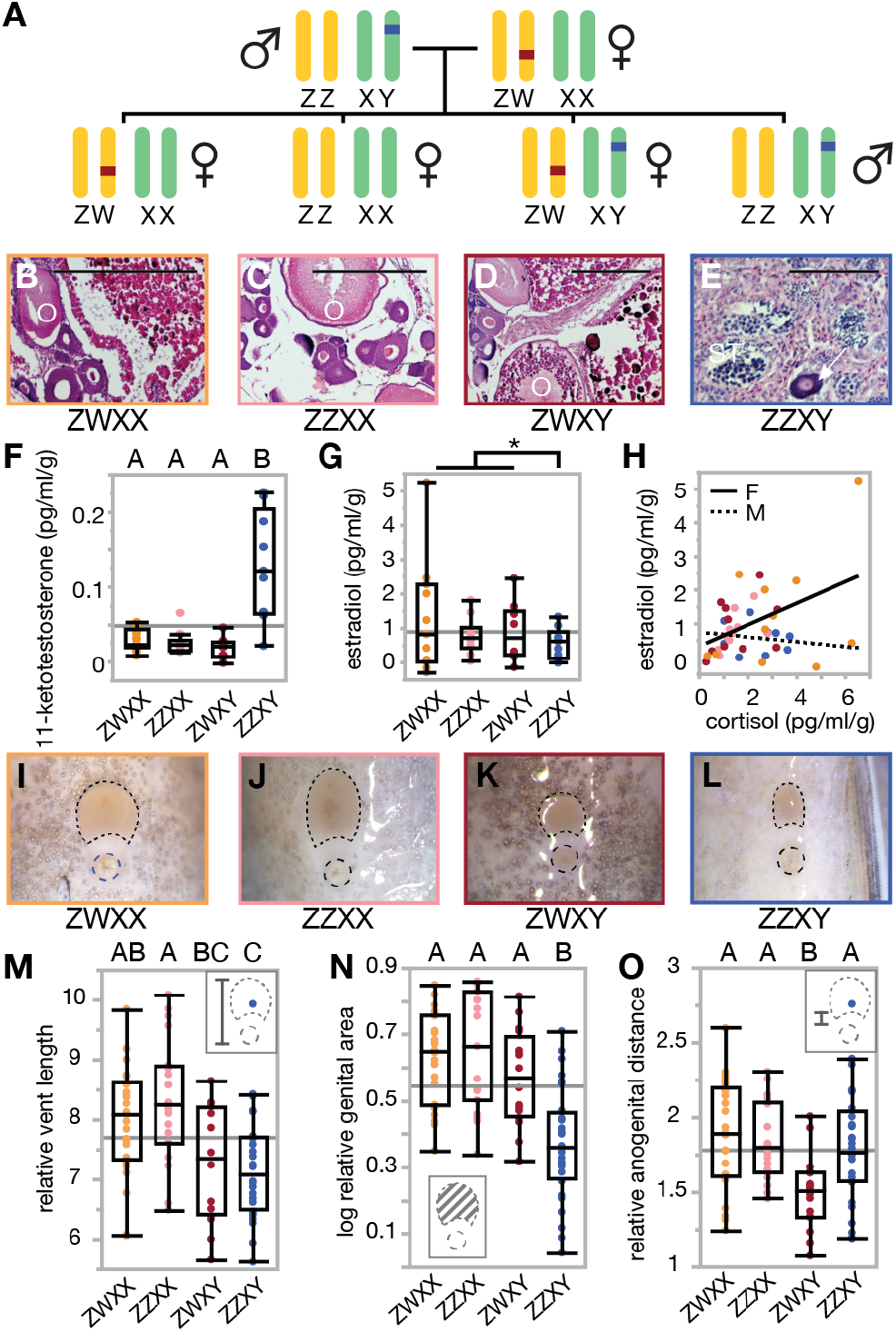
Reproductive phenotypes in a system with polygenic sex determination. Inheritance sex loci in *M. mbenjii* results in four genotypic sexes of fish, three of which are female (A). Gonads with mature oocytes (B,C,D, indicated with ‘O’) and developing sperm (E) for the sex classes support that there are two gonadal sexes, as does gonadal sex-associated levels of 11-ketotestosterone (F), estradiol (G), and cortisol association with estradiol (H). External genitalia (I, J, K, L) reflect both genotypic and gonadal sex, with vent length (M) differing by X genotype, genital area differing by gonadal sex (N), and anogenital distance modifying based on the interaction of the two (O). (histology N=3 per genotype, 12 total; hormone N=10 per genotype, 40 total; genital morphology N=139). Significance at p < 0.05 is indicated by letter grouping, from Tukey’s HSD.

Here, we look beyond color to quantify and describe the range of phenotypic variation generated among polygenic sex genotypes, and determine whether variation in traits is associated with particular sex loci or primary sex as defined by gonadal state. We find that polygenic sex genotypes impact most of the traits measured and, at the whole-organism level, produce higher-order sexual morphism than the sexual dimorphism more typically described in vertebrates. Sexual polymorphism stems from both sex-limited and sex-linked trait variation, producing broad trait variation that is complex and modular. We explore the evolutionary consequences of complex phenotypic variation associated with genotypic sexes, including how gain or loss of sex determination alleles could rapidly shift whole-organism phenotype, and how selection on specific traits could drive transitions among sex determination systems, or the maintenance of PSD systems.

## Results

### Histology, hormones, and reproductive morphology

In the wild-derived line of *Metriclima mbenjii* studied, a female with a ZW sex determiner (chr. 5) was mated to a male with an XY sex determiner (chr. 7) and produced offspring with four genotypic sexes (Fig.1A). If the two sex determiners interact additively rather than epistatically, we might predict that ZWXY females are intersex, rather than female. To test this, we examined gonad morphology, steroid hormone levels, and genital morphology. Gross examination of gonads in Y-bearing females revealed mature and developing oocytes in their gonads, and histological examination confirmed that these ovaries did not contain spermatogenic tissue. Regardless of genotypic sex, all females examined had various stages of maturing oocytes present in the gonads (Fig.1B-D), recognizable by the distinct eosin-stained (pink), round structures. In contrast, male gonads (Fig.1E) had developing sperm present in the seminiferous tubules and no mature oocytes. Of note, the male gonads had occasional immature oocytes (as indicated by an arrow), which is consistent with findings from other cichlid species both in the lab and in the wild (27) and does not appear to negatively impact reproduction for these animals. Importantly, these data confirm that ZWXY female gonads have both gross and histological features of ovaries, rather than intersex gonad tissue.

Additionally, we examined excreted levels of the primary fish estrogen (estradiol), the primary fish androgen (11-ketotestosterone), and a stress response hormone (cortisol) to determine whether ZWXY females had masculinized hormone profiles. All three female genotypes had low 11-ketotestosterone levels, feminized relative to males (ANOVA F(3,39) = 19.45, p < 0.001, Fig.1F). Estradiol results were less clear– an ANOVA with all four genotype classes did not support significant differences by genetic sex p = 0.436), however, we did find differences by gonadal sex (one-way unequal variances t-test, t = 1.97(38.93), p = 0.028, Fig.1G). Given that the three female genotypes were similar to each other, and that estradiol levels in female fishes fluctuate with reproductive cycle (28), variance among females in this measure is likely due to timing of collection with reproductive state rather than masculinization of ZWXY females. In females, cortisol had a positive linear association with estradiol levels (ANOVA F(1,30) = 7.55, p = 0.0102) while male cortisol did not (ANOVA p = 0.6737; Fig.1H). Thus, while we did not find any evidence for intrasexual variation in steroid hormones, cortisol remains a potential mediator for sex-genotype specific physiological or behavioral phenotypes, as is seen in other fishes (29).

Typically, Malawi rock cichlid males have smaller external genital openings than females, both in area and elongation (Fig.1I-L). We previously proposed that an increase in female genital size in this group has accompanied the evolution of larger eggs, and thus is functionally relevant (30). In our polygenic sex cross of *M. mbenjii*, ZWXX and ZZXX females followed this trend, as did ZZXY males. However, ZWXY females had shorter, more male-like vent lengths (full ANOVA F(3,85) = 7.12, p = 0.0003; ZWXY t = −2.42, p = 0.0179; Fig.1M), but retain a more female-like external genital area (full ANOVA F(3,85) = 17.14, p < 0.0001; ZWXY t = −1.17, p = 0.245; Fig.1N). They also possessed a shorter anogenital distance (full ANOVA F(3,85) = 4.79, p = 0.0039; ZWXY t = −3.57, p = 0.0006; Fig.1O), a trait that tends not to be predictive of sex in Malawi cichlid species, though it is a sexually dimorphic trait in other species (30, 31). In a simple sense, genital morphology of ZWXY females was intermediate to that of males and ZZXX females, reducing the overall genital area in these females. While the functional impact of this variation on reproduction is unknown, it is possible that these features may constrain egg size or influence reproductive output.

### Whole body and craniofacial morphology

Sex differences in size among fishes are common (32), which motivated us to examine whether sex genotype influences growth rate and body size. We found that growth (change in weight (g) between measurement timepoints) differed by gonadal sex, with males accumulating 0.79 ± 0.2 additional grams over a 4-month period compared to all genotypic classes of females (sex ANOVA F(1,53) = 15.35, p = 0.0003; Fig.2A). In contrast, body condition, a measure incorporating length and mass, differed by polygenic sex genotype—ZW females (ZWXX and ZWXY genotypes) had less mass per unit length than ZZ males and females (ZW ANOVA F(1,53) = 17.36, p < 0.0001; Fig.2B), suggesting ZW genotype is associated with leaner body morphology.

**Fig. 2.**
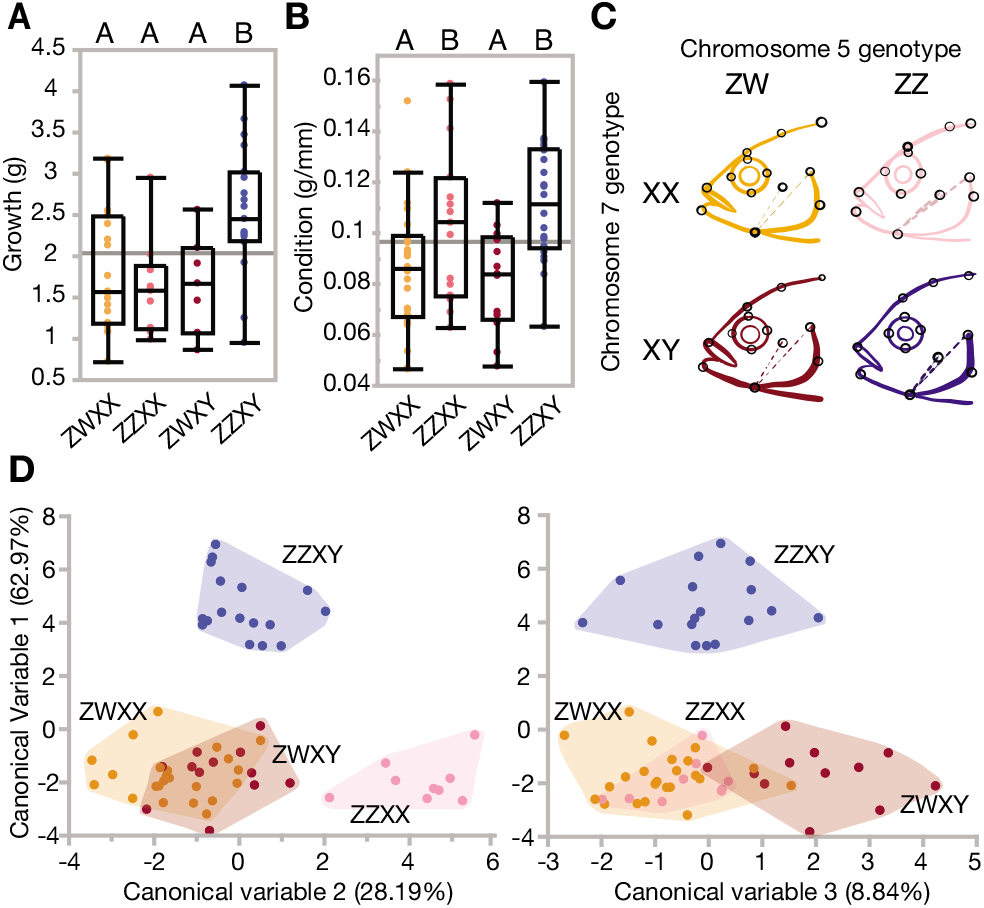
Morphology varies with polygenic sex class. Growth is higher in males (A), but condition is dependent on ZW status (B), significance at p <0.05 is indicated by letter grouping, as determined by Tukey’s HSD. Head shape is modified by both sex determiners (C)—a multivariate representation of shape quantifies differences between males and female (canonical variable 1), ZW vs ZZ females (canonical variable 2), and XX vs XY females (canonical variable 3) (D). (growth N=88, condition N =92, morphometrics N=106).

In addition to measurements of weight and length, we also evaluated body shape and size differences with morphometrics. In a model that included sex genotype, size, and sex genotype * size interaction, both size (Procrustes ANOVA; F1,53= 2.042, Z= 1.994, p < 0.05; Table S1) and sex genotype (Procrustes ANOVA; F3,53= 1.861, Z = 1.726, p<0.01; Table S1) showed significant effects on body shape. There was no significant interaction between body shape and size (Procrustes MANOVA; F3,53= 0.724, Z = 0.766, p = 0.819; Table S1), indicating the absence of allometric differences in body shape between the polygenic sex genotypes.

Previous studies indicated that sex determination loci are associated with craniofacial differences in Malawi cichlids (Parsons et al. 2015) but were unable to disentangle the effects of genetic locus and gonadal state. Here we find effects of primary sex as well as both the XY locus and the ZW locus on head shape. Canonical variate analysis (CVA) showed that ZWXX and ZZXX females have larger eyes, shallower heads, and more upturned mouths compared to ZWXY female and ZZXY male siblings (Figure 2C). Canonical variable (CV) 1 separated male fish from all female fish regardless of sex genotype (Figure 2D), and is likely mediated in part by hormones. Female siblings were distinguished on the basis of craniofacial variation by ZZ versus ZW genotype with CV2 (Figure 2D, left panel), and XX versus XY genotype with CV3 (Figure 2D), suggesting that both loci play a role in driving craniofacial variation among females. An examination of CV1 and CV3 showed similar changes to morphometric points (deeper heads, more downturned mouths, Figure S1), suggesting these differences reflect Y-specific variation, with different statistical groups indicating a sex locus by hormone interaction. A more conservative multivariate Procrustes analysis also supported an effect of genotype on craniofacial shape that was driven by both the ZW (p < 0.05) and XY (p < 0.01) loci (Table S2)

### Intestinal length

One of the most striking life history differences between male and female Malawi cichlids is that females brood their developing young in their mouths, spending weeks in a self-induced starvation state from which they readily recover once the fry are released (15). Comparisons of mouthbrooding females to those undergoing experimental starvation have shown that mouthbrooding requires a coordinated physiological response that includes adaptations of the gastrointestinal tract (21), making the gut a potential site of sexual conflict and/or sex-mediated physiology. To test for sex and sex chromosome influences on the gut, we measured intestinal length in 5-month-old *M. mbenjii* segregating both sex determination loci. Using intestinal length to test for sex-associated impacts on the gut revealed that males have the longest intestines controlling for body size (Fig. S3), and W genotype reduced intestinal length significantly (ANOVA, F(3,42) =13.08, p < 0.0001), suggesting both sex-limited and sex-linked modulation of gut length. While sex-limited differences in gut length may reflect adaptation to sexually dimorphic life history described above, implications of intrasexual variation among female genotype classes is less clear.

### Behavior

Sex differences in behavior are well documented in cichlids; in the wild, cichlid males hold territories and perform elaborate mating displays (16), while females undergo extreme behavioral shifts while mouthbrooding (33). Many species of cichlid maintain or alter their territories by shifting or moving substrate to various degrees (grooming), which is most dramatic in the species that shape elaborate bowers in the sand (34). To quantify patterns of territory use and exploratory behavior, we performed two experiments. The first was an open field assay, which measures activity, boldness, and exploration (35). We found no differences by sex genotype in movement or space use (Fig. S4). However, we found significant behavioral differences by sex in our second assay, which measured individual home tank grooming, territory occupancy and response to a novel object. In undisturbed home tanks, males spent more time out of their territories than females, and all three female genotypes behaved similarly (Fig.3A, Table S3). Males spent an estimated 35.7 seconds more time outside their territories than females (mixed model controlling for timepoint and family, sex vs null ANOVA p = 0.0002), and using genotypic sex instead of gonadal sex did not improve the model fit (p = 0.383). However, when a novel shell was introduced into the tank XY females behaved more like males (Fig.3B); spending an additional 37.6 seconds (ZWXY females) and 42.2 (ZZXY males) outside their territories than non-Y females (mixed model controlling for timepoint and family, genotype vs null ANOVA p = 0.0013; genotype vs sex ANOVA p = 0.034).

**Fig. 3.**
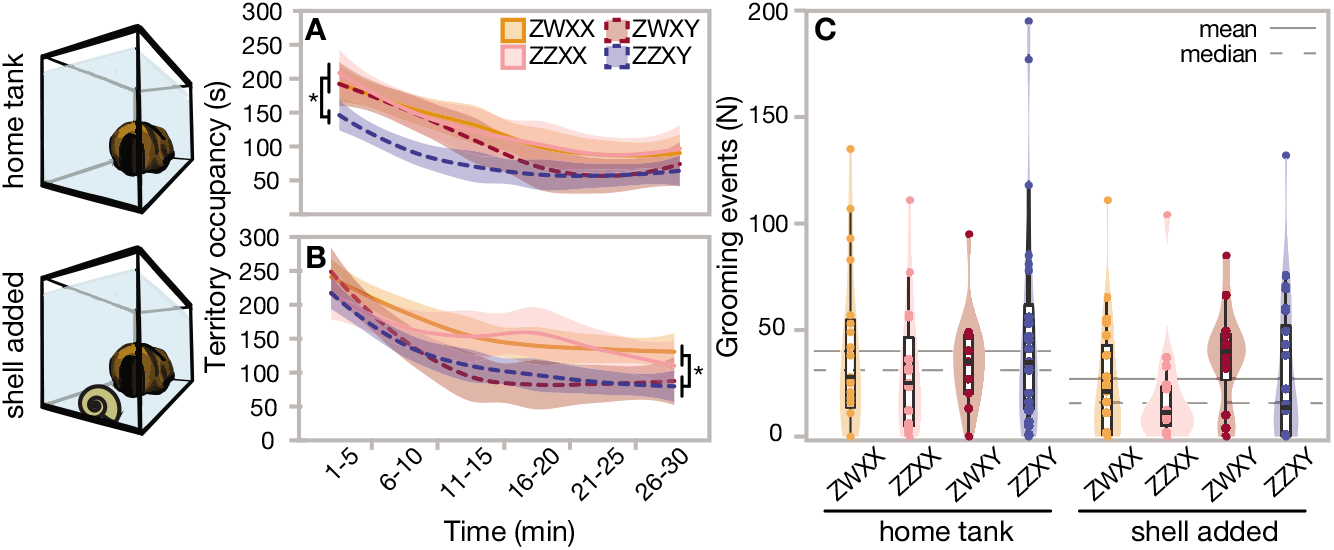
Behavioral differences among sex genotypes. XY females display more male-like behavior when confronted with novelty. There is a reduction in territory occupancy (A vs B) for XY females relative to their sisters, as well as increased grooming counts for Y-bearing individuals (C). Star indicates significance at the p < 0.05 level (N=93).

Grooming behavior was affected similarly (Fig.3C, Table S4), with Y-bearing females moving mouthfuls of sand more than their XX sisters in the face of a modified environment (Poisson general mixed model controlling for family, genotype vs null ANOVA p < 0.0001; genotype vs sex ANOVA p < 0.0001). Fish took longer to begin grooming with the addition of the shell than in the standard home tank environment (Fig. S5A; Kaplan-Meier p = 0.0047). While neither sex (Fig. S4B) nor sex genotype (Fig. S5C,D) are significant predictors of grooming latency, XY genotype has a trend towards earlier grooming (Fig. S5D, p = 0.15). These data suggest potential Y-linked effects, where XY females have bolder, more “male-like” responses to environmental disturbances. These findings are consistent with the observation that OB females in Lake Victoria are more aggressive (25), however they suggest that an increase in bold behavior is associated with the presence of the Y, rather than the presence of the W locus. Dijkstra and colleagues suggest that behavioral differences may result from modifications of the melanocortin system, which also influences pigment; if the melanocortin system were mediating this response, we would expect to see bolder behavior in the ZWXX females as well, and this is not supported by our results.

## Discussion

Here we performed phenotypic analysis of multiple traits in cichlid fish with two gonadal sexes, but four sex genotypes. The presence of multiple morphs within a given sex is not novel *per se*, and does not require PSD systems. Intriguing examples include the ruff, *Philomachus pugnax*, where variation at an autosomal locus determines three male morphs (36), and the bluehead wrasse, *Thalassoma bifasciatum*, where territorial and sneaker male morphs co-exist in a species with behavioral sex determination (37); in both species, male morphs exhibit overt differences in morphology and behavior. PSD has also been associated with variation in specific traits, including pigmentation in Lake Malawi cichlids (5) and male fitness measures in the lab in the house fly *Musca domestica* (38). The African pygmy mouse Mus minutoides presents another case of PSD impacting diverse traits. *Mus minutoides* has a single-locus sex determination system involving the common XY mammalian sex determination system, but where a novel X* chromosome variant evolved that overrides the Y, producing three female genotypes including one carrying a Y chromosome (12). Studies of *M. minutoides* indicate that Y-bearing females differ from other females in fecundity, aggression, and bite force, (39–41), indicating the presence of two multi-trait female morphs not unlike those we describe here in cichlids.

However, our findings in *M. mbenjii* present additional implications for the genetic architecture of secondary sexual characteristics. *Metriaclima mbenjii* displays intrasexual phenotypic variation associated with genotypic sex, in a diverse set of traits, including pigmentation, craniofacial and body morphology, external genital morphology, the gastrointestinal tract, and territorial behavior. Notably, no one trait is sufficient to distinguish the three female genotypes; multiple traits must be considered together to distinguish the three female classes, suggesting a greater degree of modularity of sex-associated phenotypes than other cases noted above. We propose that this modularity is due to the interplay of sex-limited and sex-linked trait variation, with multi-locus PSD systems such as the one present in *M. mbenjii* providing multiple, independently segregating sex determination loci in the genome, and therefore more possible combinations of sex-linked alleles (Fig.4). For some individual traits, Y-bearing females present more male-like or intersex-like phenotypic values, such as novel object behavior and external genital morphology, respectively. Other traits (pigmentation, body shape, and intestine length) are associated with genotype at the ZW locus. It is noteworthy that at least for one trait, multiple W-linked alleles have evolved that produce additional variation within a W-linked pigmentation phenotype (26), demonstrating that sex-linked variation is not limited to two alleles. Modularity due to the interaction of sex-limitation and sex-linkage is further supported by multivariate analysis of craniofacial morphology, where sex-limited variation appears to drive the first canonical variable, and sex-linked variation at both the XY and ZW loci drive differences in the second and third canonical variables in females. The genetic architecture of secondary sexual characteristics in *M. mbenjii* presents interesting evolutionary implications. Surveys of sex determination systems among cichlids suggest that the chr. 5 ZW system is recently-derived relative to the more widespread and common chr. 7 XY system (11, 42). Considering the chr. 5 W allele as a relatively recent introduction to some species suggests additional context for our results. We previously showed that the W haplotype carries a dominant derived allele of *pax7a* that produces the orange blotch color morph (5, 26); here we show that the haplotype is also associated with variation in other traits and can shift variation in morphological traits into new phenotypic space, producing more slender body morphology, shorter relative gut length, and novel aspects of craniofacial variation not present in males or females of a purely XY population. That is, the presence of the ZW system does not simply produce intermediate secondary sexual characteristics in individuals carrying both a W and a Y allele, instead the addition of the W produces phenotypic novelty within traits, as well as new modular combinations of traits not present in the ancestral population.

**Fig. 4.**
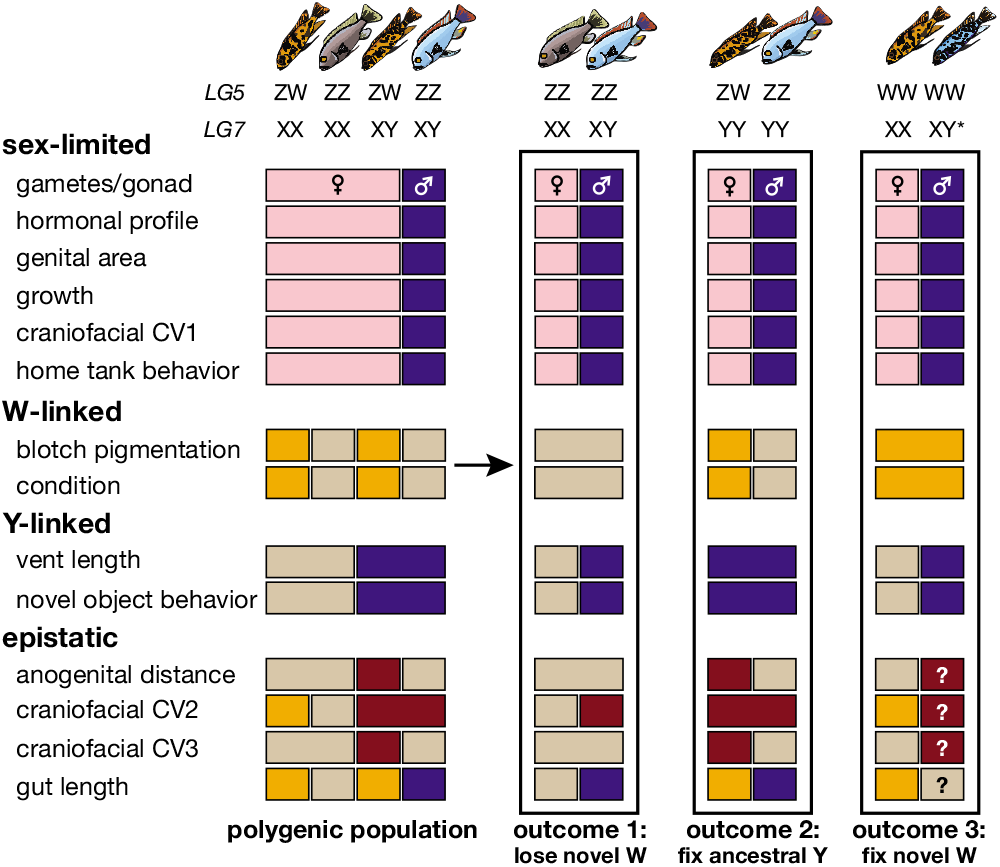
Modular phenotypic variation associated with sex genotypes in *M. mbenjii*. Left side, graphical summary of phenotype differences in population with PSD, sorted by inferred cause of variation. Right side, predicted evolutionary outcomes of loss or fixation of sex determination alleles. Different colors indicate statistically distinct groups. In outcome 3, the star next to the Y indicates a novel Y allele epistatically insensitive to the W.

Comparative surveys also indicate that either the chr. 5 ZW or chr. 7 XY system on its own is sufficient to determine primary sex for a species (5, 11), suggesting that dominant sex determination alleles can be lost or fixed at one locus without impacting sexual development. Where they co-occur in pure species, the chr. 5 W allele epistatically dominates the XY system, such that individuals with both a W and a Y develop as females, and males with the W-linked blotch color morph are rare in natural populations (16, 43). However, it is possible that genetic background or allelic variation at sex alleles can shift the epistatic hierarchy; in-lab interspecific crosses between a ZWXX female of *M. mbenjii* and a ZZXY male of *Aulonocara koningsi* show the chr. 7 XY controlling sex despite the presence of a chr. 5 ZW (Supplementary Table S5). Thus, the hierarchy of sex determination systems can also seemingly shift with differences in genetic background. Despite strong pre-mating isolation in the wild, occasional hybridization provides opportunity for such changes in genetic background, and allows for introgression of genetic variation among cichlid species (44). Further, introgression of sex determination alleles across genera is supported by analysis of the chr. 5 sex determination locus, where a W-specific haplotype indicates a single origin of the W allele that is now found in dozens of species from four genera (5, 26). We note that a supernumerary B chromosome associated with female sex determination or sex reversal in at least three genera of Malawi cichlid has been reported at low frequency in *M. mbenjii* (1 in 60 individuals) (45, 46); while the B chromosome is not present in families in this study, in wild populations such low frequency sex determining alleles may be available for selection to act upon to shift whole organism phenotypes or sex ratio.

Thus, there is demonstrated opportunity for relatively common evolutionary gains and losses of genetic sex determination systems among Lake Malawi cichlids, with concomitant shifts in any traits impacted by sex-linked alleles at those loci. More broadly stated, evolutionary transitions among genetic sex determination systems have impacts beyond the gonad, with potential to remodel the whole organism in unexpected ways (Fig.4). Modular whole-organism phenotypes arising from the interplay of sex-limited and sex-linked traits in cichlids with PSD may produce alternative fitness strategies and be stably maintained by selection (47–49). We consider variation in *M. mbenjii* by sex genotype in light of these hypotheses Fig.4. If color or behavior differences provide alternative predator avoidance tactics, or head shape or gastrointestinal differences support alternative foraging specializations, balancing selection may maintain multiple morphs and their associated sex determination loci. In the case of rock-dwelling cichlids, this balancing selection may arise from high competition in a structurally, trophically, and visually complex environment, or through temporal differences in substrate appearance and food availability over seasons or years (47). On the other hand, if directional selection favors certain trait values or combinations of traits present in one morph, those trait values may arise in a population by loss or fixation of various sex determination alleles Fig.4.

Alternatively, if a particular sex determination system confers an advantage under certain conditions, it may rise in frequency and drive shifts in a number of associated traits. For example, if the ancestral Y became fixed in our model population 4, the population would rely on the ZW locus for sex determination and females would display a subset of phenotypic variation found in the ancestral population. The result would be females displaying only blotched pigmentation, slender body morphology, more male-like territorial behavior, intermediate external genital morphology, and shallower heads (ZW) with deeper eyes and more downturned mouths (XY) that reflect a shift in sex-specific craniofacial variation. Thus, while directional selection on a single trait in one or both sexes could drive transitions between genetic sex determination systems as previously hypothesized (5, 49), our results here suggest selection on a single trait or sex determination locus could shift an array of sex-linked and sex-limited phenotypes to rapidly and flexibly change whole-organism phenotypes. Due to the modular nature of sex-limited and sex-linked trait variation in polygenetic sex determination systems, selection on different aspects of an organism could drive a number of evolutionary trajectories in a population. While we suggest adaptive hypotheses above for trait variation and population evolution, non-adaptive scenarios are also likely, including drift and the accumulation of near-neutral allelic variation at sex determination loci.

Longstanding questions in biology surround what factors support or drive diversity of species and sex determination systems. African cichlids serve as model radiation exhibiting extremely high diversity of both, providing the possibility that rapid diversification of genetic sex determination systems may support rapid speciation, or vice versa. Here, we suggest that the modular, higher-order sex polymorphism that can be produced by PSD systems has the potential for profound evolutionary consequences. The increased phenotypic variation surrounding whole organism phenotypes and ease with which sex-specificity can be shifted may be responsible, in part, for the extreme parallel diversity found in phenotypes, sex determination systems, and species in the explosive radiation of African cichlid fishes. Ultimately, detailed genetic and phenotypic analysis within and among natural cichlid species populations are necessary to determine how variation in sex determination systems impacts organismal traits and fitness through space and time.

## Materials and Methods

### Animals and genotyping

A wild-derived line of *Metriaclima mbenjii* was maintained under Institutional Animal Care and Use Committee (IACUC) guidelines (Protocols 17-101-O and 17-140-O). We established a breeding group with an OB female (ZWXX) and dominant male (ZZXY). Offspring were collected from F1 and F2 (ZWXX x ZZXY) sibling matings. We identified chr. 5 ZW genotype by eye because it is linked to the OB pigmentation polymorphism. We genotyped the male (XY) system on chr. 7 using established markers (11, 42).

### Tagging and gross measurements

All individuals were anesthetized and measured for standard length (mm) and weight (g), injected with a passive integrated transponder (PIT) radio tag, and had fin tissue collected for genotyping. Following final assays, whole fish ventral photographs and high-definition head photographs were taken using a dissecting microscope, and an additional round of weight and standard-length measurements were taken.

### Behavioral Assays

For the open field assay, fish were netted from home tanks and placed into a square arena for five minutes, and tracked using C-Trax (v0.5.4) (50). Summary values for position and speed in the arena were generated using custom R-scripts (R v3.3.1; (35)). For the home tank behavior and novel object assays, cohorts of up to ten fish from all sex genotypes were acclimated to individual arena tanks for 24 hours. They were then filmed for two 30-minute assays, once without disturbance and once after adding a novel object (shell). The videos were evaluated for territory occupancy and grooming instances using the Observational Data Recorder (ODRec v2.0 beta), and summary statistics were generated with custom R-scripts.

### Hormones

We used a non-lethal method for collecting hormones from holding water (51) and quantified 40 individuals for cortisol, 11-ketotestosterone, and estradiol using a colorimetric enzyme-linked immunoassay following manufacturer’s instructions (EIA, Cayman chemicals).

### Gut Length

To adequately control for plasticity in gastrointestinal development, we raised two families in a separate cohort for gut length measurements. Controlling for density and food amount, fish were humanely euthanized with 250 mg/L tricaine methanesulfonate (MS-222) at 5 months old and measured for standard length, weight, and gut length.

### Gonad histology

Once all assays were complete for a family, fish were humanely euthanized with 250 mg/L MS-222. Gonads were extracted and preserved in 10% neutral-buffered formalin, and processed for histological staining by wax embedding. Slides were stained with hematoxylin and eosin (HI&E) to visualize gonadal tissue features.

### Morphometrics and statistics

From photographs, 20 landmarks (Fig.S1) were digitized for geometric morphometric analyses using the R package geomorph (52). Landmarks were superimposed by a Generalized Procrustes Analysis (53), and genotype effects were modeled using log-transformed centroid size as a covariate to control for allometry (shape genotype * size). Additionally, we used a Procrustes ANOVA with permutation (52) to quantify shape variation associated with sex genotype, with differences between genotype identified with post-hoc pairwise comparisons (54). Anogenital measurements were quantified from ventral photographs as described previously(30).

All other statistical analyses were performed using packages available in JMP v12 (SAS) and R v3.6.1(55). For genital morphology, hormones, and gut length, sex genotype effects were modeled using ANOVA and pairwise differences tested with Tukey’s honestly significant difference (HSD) test (JMP). Gut length increases allometrically with body size, so we modeled genotype after controlling for log10 mass. For behavioral tests, the amount of time spent in the cave territory was modeled with a mixed model including genotype and time as fixed effects and family as a random effect (56). Counts of grooming, controlling for family as a random effect, were modeled with a Poisson generalized regression (57). Latency to groom was modeled with a survival model (58). Values reported with ± SEM.

## ACKNOWLEDGMENTS

The work was funded by support to RBR from the National Science Foundation (IOS-1456765), an Arnold and Mabel Beckman Institute Young Investigator Award to RBR, a Keck Center for Behavioral Biology Award to ECM, and an NCSU Provost Award to ECM. Thanks to M. Kaitlin Barker and Lynea Bull for assistance with genotyping families, Natalie Roberts and Kate Coyle for assistance with fish husbandry, and Matt Bertone for assistance with building the light box for photography.

---

**Fig. S1.**
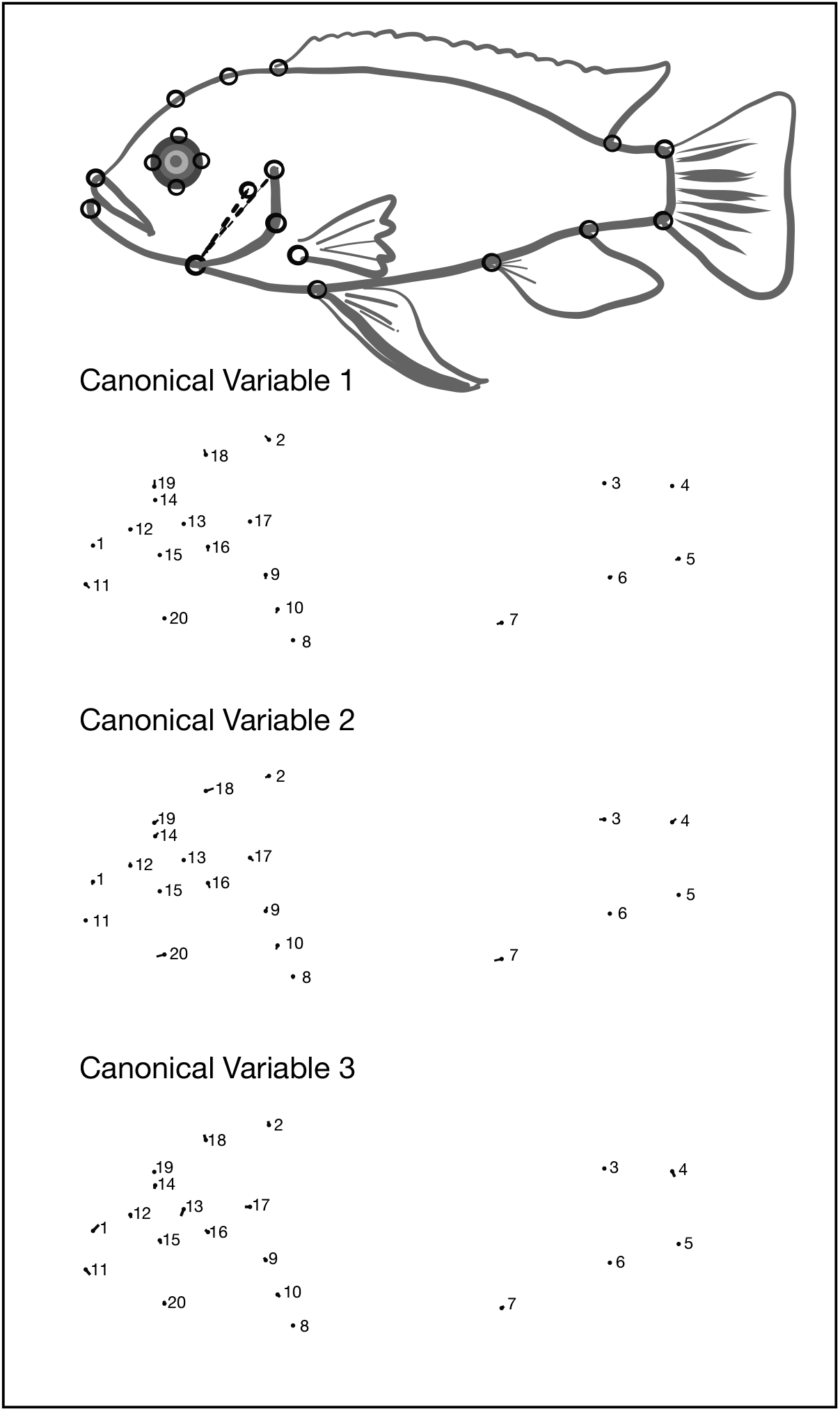
Depictions of which morphometric landmark shifts comprise each canonical variable. Each dot indicates a standard position of a morphometric landmark when all groups are considered; all vectors directed off the landmarks indicate the specific shape change described by the multivariate canonical variable. Numbered points are as follows: 1) anterior tip of maxilla; 2) anterior insertion of dorsal fin; 3) posterior insertion of dorsal fin; 4) dorsal insertion of caudal fin; 5) ventral insertion of caudal fin; 6) posterior insertion of anal fin; 7) anterior insertion of anal fin; 8) insertion of pelvic fin; 9) dorsal insertion of pectoral fin; 10) ventral insertion of pectoral fin; 11) anterior tip of dentary; 12) anterior-most edge of eye; 13) posterior-most edge of eye; 14) dorsal-most edge of the eye; 15) ventral-most edge of eye; 16) dorsal tip of preoperculum; 17) dorsal tip of operculum; 18) intersection of the dorsal tip of the preoperculum with the dorsal profile of the body; 19) intersection of the dorsal-most edge of eye with the dorsal profile of the body; 20) intersection of the ventral-most edge eye with the ventral profile of the body

**Fig. S2.**
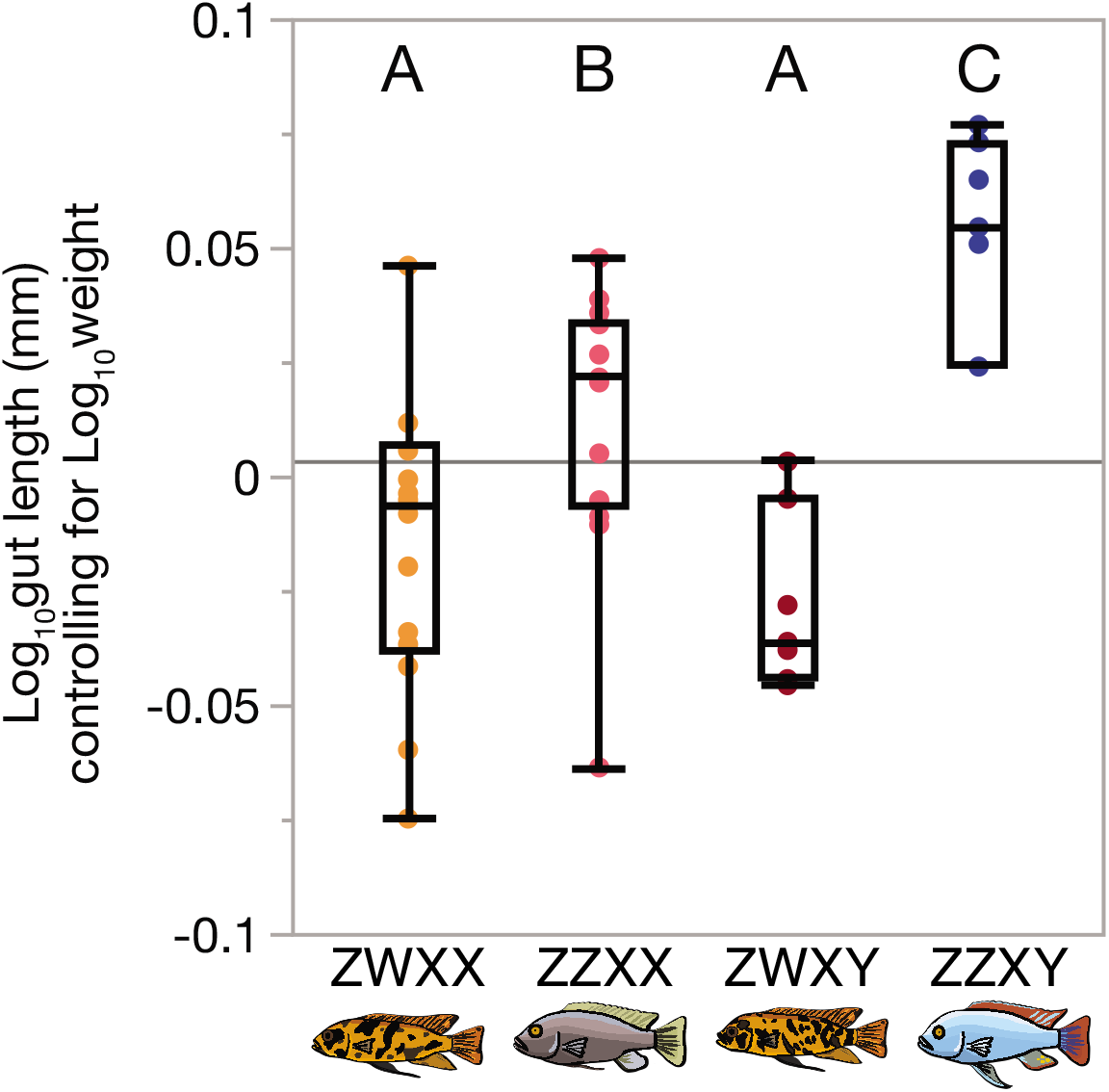
Gut length differs by genotypic sex class. Residuals from a model where Log10 Gut Length (mm) Log10 weight show differences by sex genotype. Letters indicate statistical grouping determined by Tukey’s HSD, p > 0.05

**Fig. S3.**
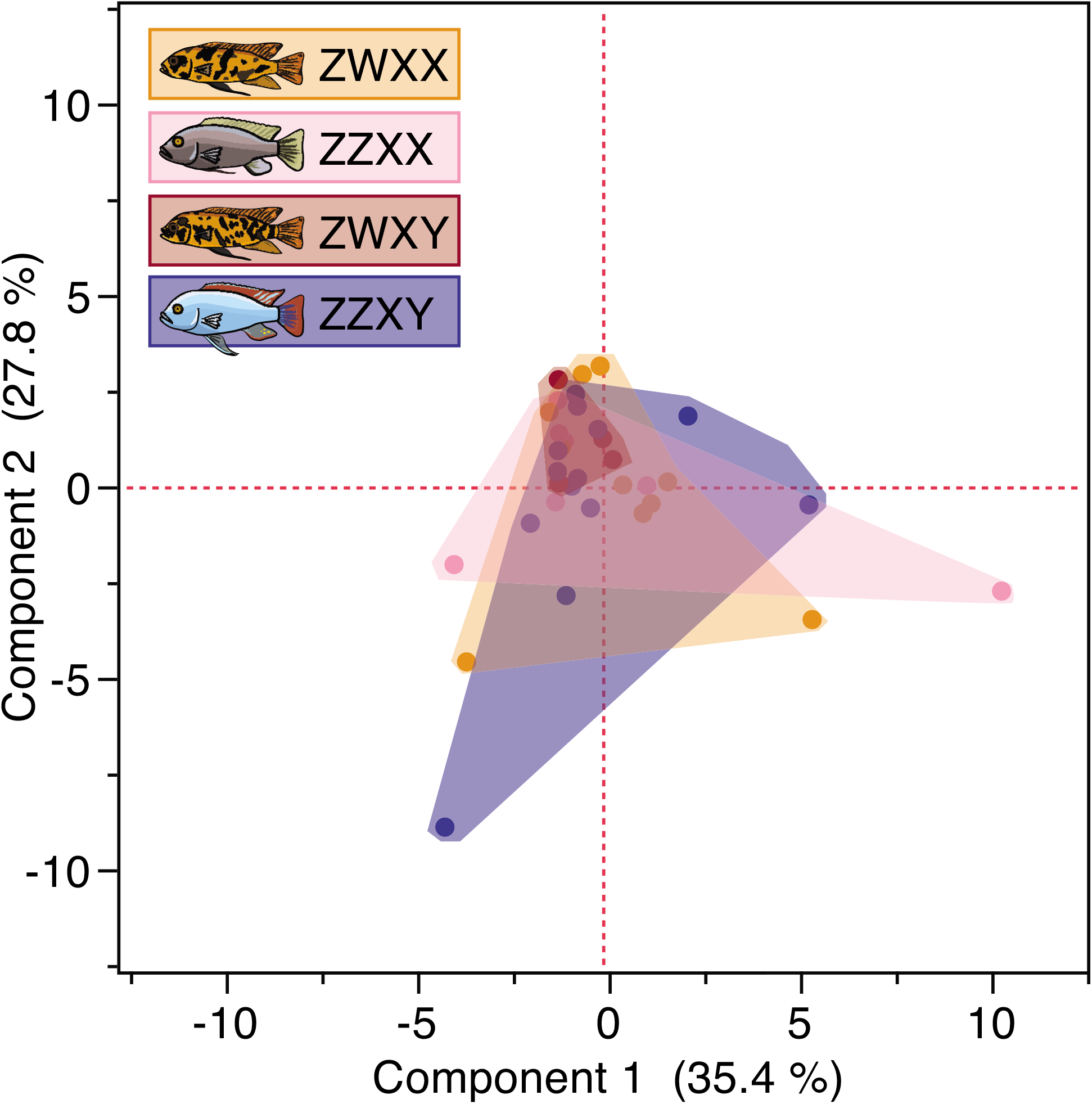
Open field behaviors are not different by sex, as indicated by overlapping principle components 1 and 2 by genotype class

**Fig. S4.**
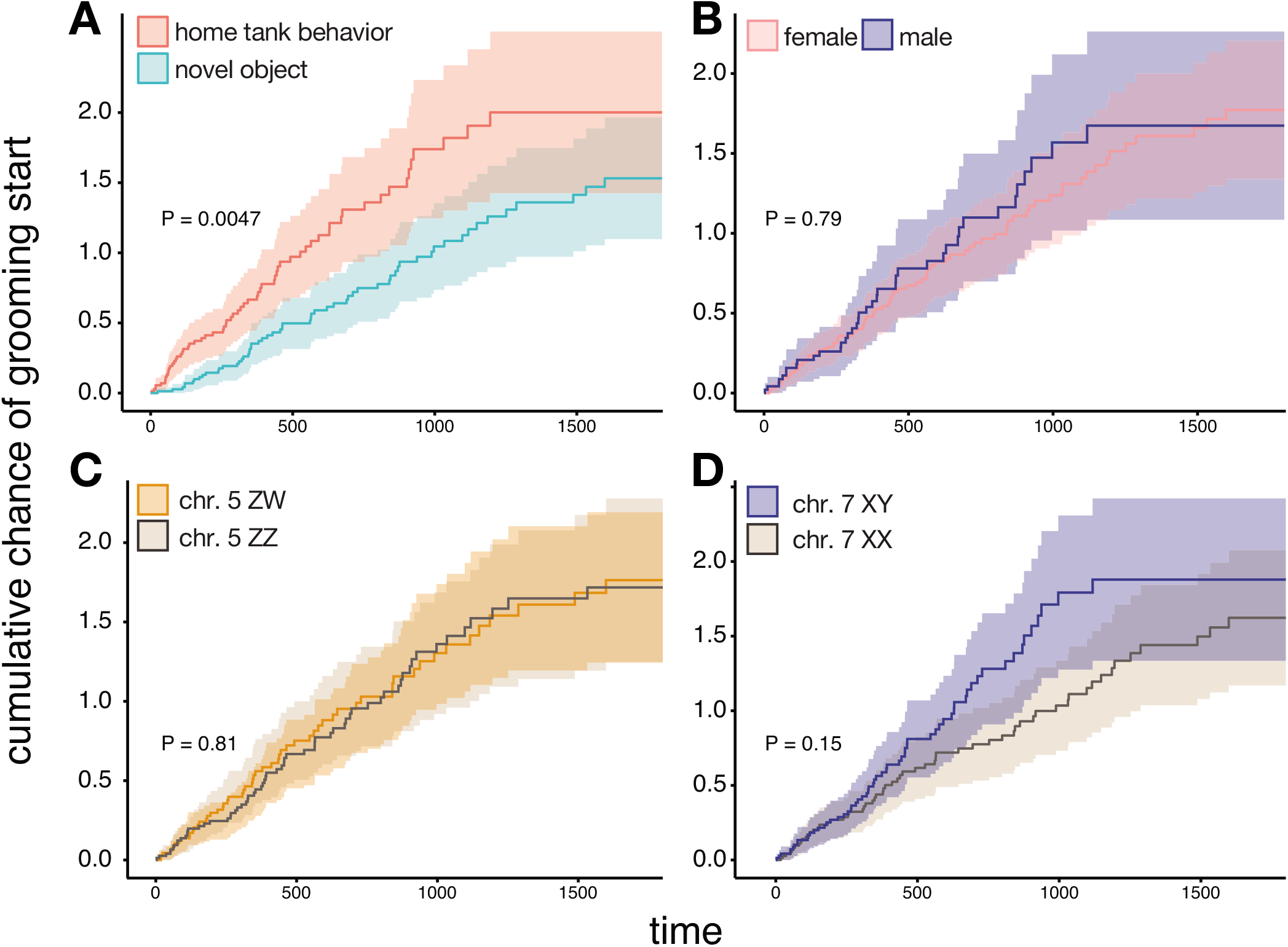
Grooming latency. Cumulative hazard plots show the chance over time that individuals resumed territory grooming activity after the start of filming. P-values in panel result from Kaplan-Meier models of latency, with the assay (A), gonadal sex (B), chr. 5 sex genotype (C), and chr. 7 sex genotype (D) as model main effects

**Table S1.** Procrustes ANOVA of body shape and polygenic sex genotype. P values are from random residual permutation procedures (n = 1000). Significant effects in bold.

|  | df | SS | MS | F | P |
| --- | --- | --- | --- | --- | --- |
| Genotype | 3 | 0.003 | 0.001 | 1.861 | **0.004** |
| Log centroid size | 1 | 0.001 | 0.001 | 2.042 | **0.022** |
| Genotype*Log centroid size | 3 | 0.001 | 0.0003 | 0.724 | 0.819 |
| Residuals | 53 | 0.028 | 0.001 |  |  |
| Total | 60 | 0.034 |  |  |  |

**Table S2.** Multivariate Procrustes distance metric by sex genotype.

| Distances among groups |  |  |  |
| --- | --- | --- | --- |
|  | ZWXX | ZWXY | ZZXX |
| ZWXY | 0.0062 | - | - |
| ZZXX | 0.0127 | 0.0131 | - |
| ZZXY | 0.0127 | 0.0113 | 0.014 |
| P-values from permutation tests (10000 rounds) |  |  |  |
|  | ZWXX | ZWXY | ZZXX |
| ZWXY | 0.8876 | - | - |
| ZZXX | **0.0272** | 0.1259 | - |
| ZZXY | **0.0015** | 0.0924 | **0.0204** |

**Table S3.** Territory occupancy models.

|  | home tank |  |  | shell |  |  |
| --- | --- | --- | --- | --- | --- | --- |
| Sex vs null | **P = 0.0002069** |  |  | **P = 0.00286** |  |  |
| Genotype vs null | **P = 0.001314** |  |  | **P = 0.001323** |  |  |
| Genotype vs sex | P = 0.3827 |  |  | **P = 0.03374** |  |  |
|  | estimate | Z val | Pval | estimate | Z val | Pval |
| ZWXY | -13.327 ± 13.8 | -0.965 | 0.33484 | -37.5779 ± 14.8 | -2.538 | **0.011487** |
| ZZXX | 7.059 ± 12.8 | 0.551 | 0.548819 | -8.2461 ± 13.7 | -0.6 | 0.548819 |
| ZZXY | -36.669 ± 11.3 | -3.254 | **0.00123** | -42.1775 ± 12.1 | -3.499 | **0.000513** |
| time | -4.087 ± 0.52 | -7.83 | **3.73E-14** | -4.6171 ± 0.56 | -8.211 | **2.36E-15** |

**Table S4.**
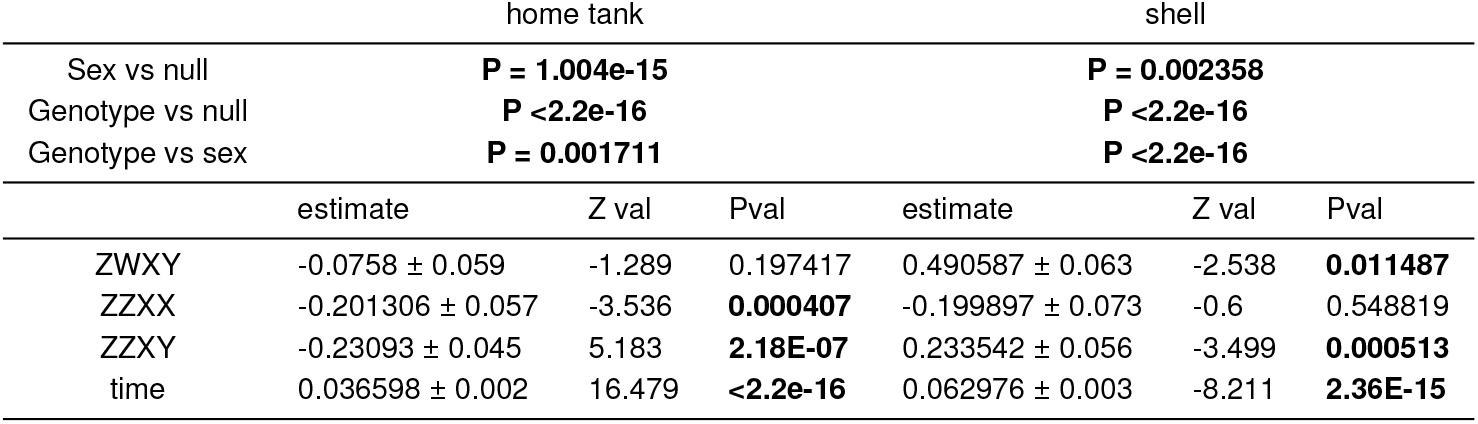
Grooming count models.

**Table S5.** Primary sex by sex genotype in Metriaclima mbenjii x Aulonocara koningsi F2 hybrids, from a cross of ZWXX x ZZXY F1 hybrids.

| Genotype | Female | Male |
| --- | --- | --- |
| ZZXX | 41 | 0 |
| ZWXX | 44 | 0 |
| ZWXY | 0 | 35 |
| ZZXY | 0 | 47 |

